# Time-dependent transcriptomic changes following protoplast isolation in plants

**DOI:** 10.64898/2026.07.14.738454

**Authors:** Hao Zhang, Ankush Sangra, Anita Giabardo, Joshua C. Wood, Julia Brose, Sierra S. Cloud, John P. Hamilton, Kathrine Mailloux, Brieanne Vaillancourt, C. Robin Buell, Robert J. Schmitz

## Abstract

Protoplast isolation is widely used for plant functional genomics and single-cell analyses, but its impact on transcriptional and cell state dynamics remains incompletely understood. Here, we generated time-course RNA-seq data from leaf protoplasts of *Arabidopsis*, maize, and poplar, sampling at multiple time points following isolation, to systematically characterize global transcriptional dynamics across species. We identified two major drivers of transcriptional variation: a persistent protoplast isolation effect and a progressive time-dependent transcriptional program, which can be divided into early, middle, and late stages corresponding to an immediate stress response, metabolic and chromatin regulation dynamics, and sustained metabolic and proteostasis regulation, together with species-specific differences across stages. We observed a rapid loss of cell-type-specific transcriptional signatures within 6 hours in *Arabidopsis* and maize, whereas poplar showed a slower decline. Single-nucleus RNA-seq at 6 hours in maize confirmed attenuation of cell-type-specific transcriptional structure. Furthermore, leveraging this time-course dataset enables the identification of aberrant cell states in single-cell RNA-seq data, exemplified by clusters showing elevated activity of protoplast isolation-associated, middle-, and late-stage transcriptional programs characteristic of stress-like states. Together, our results provide a cross-species framework for dissecting protoplast-induced transcriptional and cell state dynamics and facilitate the systematic identification of stress-associated cell states in single-cell transcriptomic data.

## Introduction

Protoplasts, generated by enzymatic removal of the cell wall, provide a versatile system for transient gene expression (Chen *et al*., 2023; Miao and Jiang, 2007), genome editing (Hsieh *et al*., 2026; Yue *et al*., 2021), *in vitro* culture (Pasternak *et al*., 2020; Yoo *et al*., 2007), and have long served as a model for investigating dedifferentiation and regeneration (Davey *et al*., 2005; Li *et al*., 2024a; Reed and Bargmann, 2021). In parallel, protoplast isolation is widely used in plant single-cell transcriptomic workflows, which have enabled high-resolution characterization of cell identities and their dynamic changes, systematic definition of cell types across diverse tissues and species, reconstruction of developmental trajectories, and identification of gene regulatory networks underlying cell fate specification, differentiation, and state transitions (Denyer and Timmermans, 2022; Li *et al*., 2016; Procko *et al*., 2022; Shahan *et al*., 2022; Shulse *et al*., 2019; Yin *et al*., 2024; Zheng *et al*., 2023). Together, these applications underscore the broad utility and importance of protoplasts in plant cell biology and functional genomics.

Emerging evidence suggests that protoplast isolation is not a passive sampling procedure, but rather a procedure that triggers rapid and widespread transcriptional and chromatin-level dynamics (Chupeau *et al*., 2013; Liu *et al*., 2022; Xu *et al*., 2021a), which including immediate activation of stress-responsive pathways (Král *et al*., 2025; Li *et al*., 2024b). These responses indicate that removal of the cell wall and dissociation from native tissue architecture triggers disruption of cellular homeostasis, and chromatin-associated regulatory states. In addition, protoplast isolation alters metabolic activity and energy allocation, reflecting a shift from tissue-integrated function to a stress-adaptive cellular state (Mukundan *et al*., 2025; Ondřej *et al*., 2010; Pasternak *et al*., 2020; Zhang *et al*., 2022). Despite these observations, the most current understanding is based on endpoint comparisons, leaving the early temporal dynamics of protoplast-induced transcriptional changes poorly resolved, including whether transcriptional programs remain stable or undergo time-dependent changes following isolation and the kinetics of such changes.

Here, we generated time-course RNA-seq datasets spanning different protoplast incubation time points from leaf cells from *Arabidopsis*, maize, and poplar to systematically characterize the dynamics of transcriptional responses to cellular isolation and transcriptional changes. We found a persistent protoplast isolation effect and a progressive time-dependent transcriptional programs across three species, affecting distinct functional pathways. By single-nucleus RNA-seq (snRNA-seq), we show that protoplast isolation triggers rapid erosion of differentiated cell identity, with a marked attenuation of cell type–specific transcriptional signatures within six hours in maize. Thus, our findings provide a cross-species framework for understanding how cellular isolation reshapes plant transcriptional programs and progressively erodes differentiated cell-type–specific signatures.

## Results

### Global transcriptional dynamics of protoplasts reveal isolation and time effects across species

To comprehensively understand the dynamics of isolated protoplasts, we generated time-course RNA-seq libraries after leaf protoplast isolation in three species: *Arabidopsis thaliana* (*Arabidopsis*), *Zea mays* (maize), and *P. tremula* x *P. alba* INRA 717-1B4 (poplar) (**Fig. 1A; Supplementary Table S1**). We then quantified global transcriptomic distances between protoplasts at different time points and leaf tissue, revealing two major factors influencing protoplast dynamics in plants: the protoplast isolation effect, in which isolated protoplasts exhibit strong transcriptomic divergence from leaf cells, and the time effect, during which protoplasts gradually change over time (**Fig. 1B**). Hierarchical clustering and correlation analysis of the whole transcriptome grouped protoplasts across time points into three stages. Here, 0 h denotes freshly isolated protoplasts collected immediately after 2 h of cell wall digestion. The early stage comprised 0 and 3 h in Arabidopsis and maize and 0 h in poplar; the middle stage comprised 6 and 9 h in Arabidopsis and maize and 3 and 6 h in poplar; and the late stage comprised 16–24 h (**Fig. S1A**).

**Figure 1.**
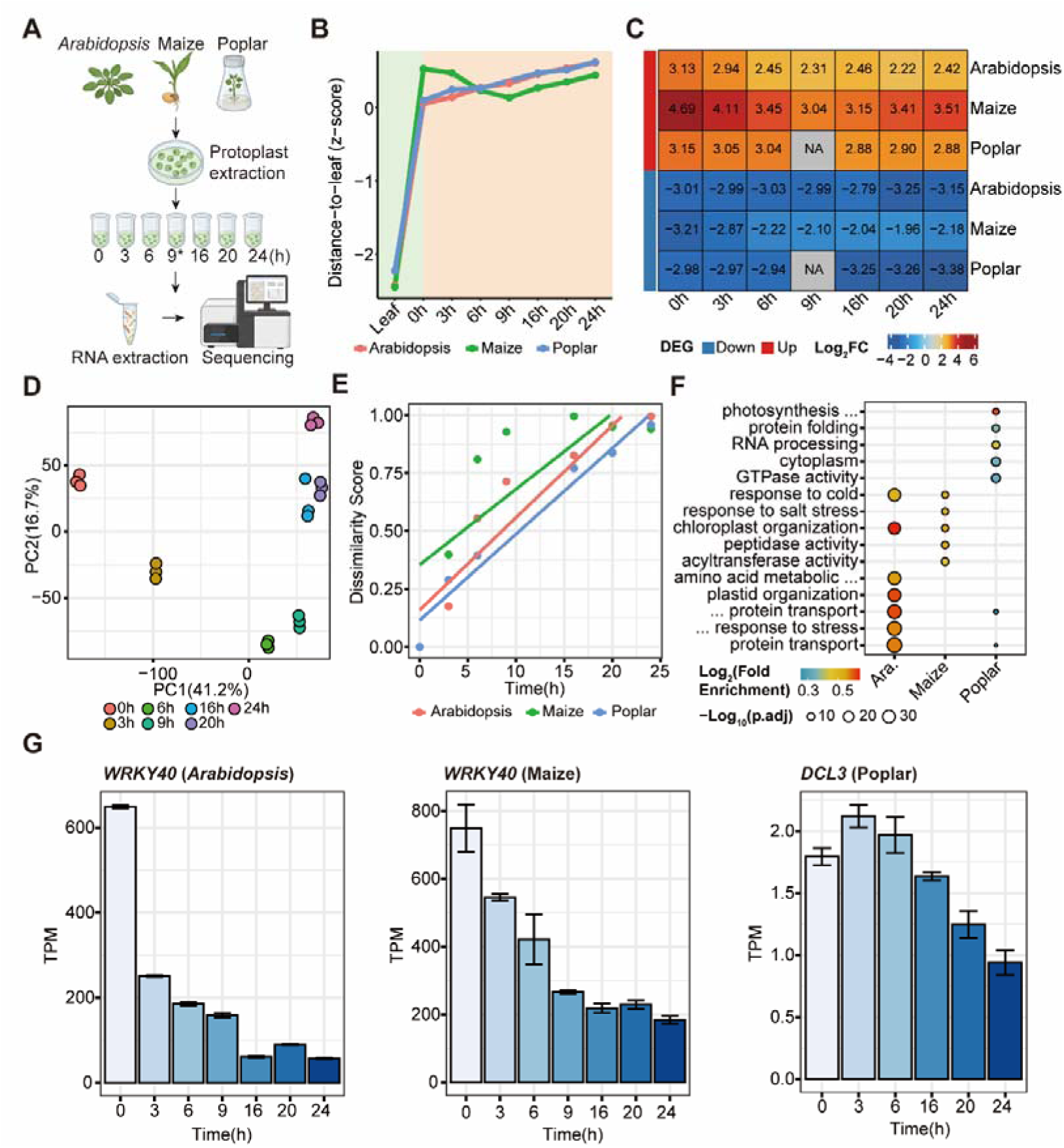
Global transcriptional dynamics of protoplasts reveal isolation and time effects. **A.** Schematic overview of the experimental design showing leaf protoplast isolation and time-course RNA-seq sampling across *Arabidopsis*, maize, and poplar. *9 h was not sampled in poplar. **B.** Line plot showing global transcriptomic distance to leaf tissue across three species. **C.** Heatmap showing fold changes of isolation-induced DEGs relative to leaf tissue across three species. **D.** PCA of gene expression at different time points in *Arabidopsis*. **E.** Linear regression of dissimilarity score against time across three species. **F.** GO dot plot showing functional enrichment of genes significantly associated with time across three species. **G.** Expression patterns of *WRKY40* in Arabidopsis and maize, and *DCL3* in poplar.

To uncover protoplast isolation effect, we identified differentially expressed genes (DEGs) between leaf tissue and fresh protoplasts (0 h) in all three species (**Fig. S1B**). Functional enrichment analysis showed that up- and down-regulated DEGs were associated with distinct biological processes: stress-response pathways (e.g., *JAZ1*; **Fig. S1B**) were consistently up-regulated, whereas photosynthesis-related (e.g., *CRY1*; **Fig. S1B**) and cell wall-related pathways (e.g., *CESA3*, *CESA4*, and *CESA5*; **Fig. S1B**) were consistently down-regulated across all three species (**Fig. S1C-E**). Notably, the isolation-associated expression patterns were largely maintained across subsequent time points, as up-regulated genes consistently showed positive fold changes, whereas down-regulated genes remained negatively correlated across time points in all three species (**Fig. 1C**). Collectively, these results indicate a robust isolation-induced transcriptional effect that persists across time and species.

Principal component analysis (PCA) revealed a gradual shift in transcriptomic profiles over time across the three species (**Fig.1D**, **Fig. S2A**). We defined a dissimilarity score and scaled it to a 0–1 range to quantify the effect of time on transcriptomic divergence (1 − similarity to the 0 h sample). Linear regression revealed a strong linear relationship between the dissimilarity score and time, with high R² values across *Arabidopsis* (0.872), maize (0.606), and poplar (0.952) (**Fig.1E**, **Fig. S2B**). Collectively, these results indicate that isolated protoplasts undergo a continuous, time-driven transcriptional reconfiguration process. Moreover, quantification of transcriptome dissimilarity provides an opportunity to pinpoint transcriptional signatures underlying the time effect. We identified a large number of genes (75.6% in *Arabidopsis*, 46.2% in maize, and 61.8% in poplar) that were significantly associated with time across the three species using a time-course modeling approach (FDR < 0.05; **Fig. S2C**). The top associated Gene Ontology (GO) terms differed among the three species. Arabidopsis and maize were mainly associated with stress-related responses, including *WRKY40* in both species (**Fig. 1G**). In addition, maize showed enrichment in enzymatic activity–related terms and protein transport, whereas Arabidopsis was enriched in amino acid metabolism. In contrast, poplar was enriched in fundamental cellular processes such as protein folding and RNA processing, including *DCL3* (**Fig. 1F**, **1G**). To evaluate the potential contribution of circadian regulation, we analyzed the expression of canonical circadian clock genes across the three species. These genes exhibited rhythmic oscillations throughout the time course (**Fig. S3A-C**), indicating the presence of an active circadian program. In contrast, the time-associated genes exhibited non-oscillatory expression patterns (**Fig. 1G**), suggesting that their transcriptional changes are unlikely to be primarily driven by circadian regulation. Overall, time exerts a strong influence on the global transcriptional dynamics of isolated protoplasts across all three species.

### Temporal transcriptional programs during protoplast incubation across species

Using non-negative matrix factorization, we decomposed transcriptomic profiles into three major programs across the time course, identifying early, middle, and late programs that were consistently recovered across all three species, in agreement with results from hierarchical clustering and correlation analysis (**Fig. 2A**). To characterize the functional properties of these programs, we performed GO enrichment analysis using the top-ranked genes associated with each program in each species, with the top 100 genes used for visualization as similar enrichment patterns were observed using an alternative cutoff of 500 genes (**Fig. S4A**). The early program was characterized by rapid recovery of cellular activity, including reactivation of photosynthesis, translational initiation, protein folding, and redox homeostasis, together with responses to light and oxidative stress (**Fig. S4B**). The middle program reflected a transitional state dominated by metabolic reprogramming and hormone-mediated regulation, including carbohydrate metabolism, glycolysis, respiration, and responses to cytokinin, gibberellin, and nutrient availability (**Fig. S4C**). In contrast, the late program was enriched for stress defense and immune-related processes, including responses to wounding, fungal infection, jasmonic acid, and oxidative stress, as well as protein degradation-related activities and small-molecule catabolism (**Fig. S4D**). The early, middle, and late programs define a sequential transcriptional cascade from cellular recovery to metabolic and hormonal reprogramming, and ultimately to sustained stress and defense responses during protoplast incubation.

**Figure 2.**
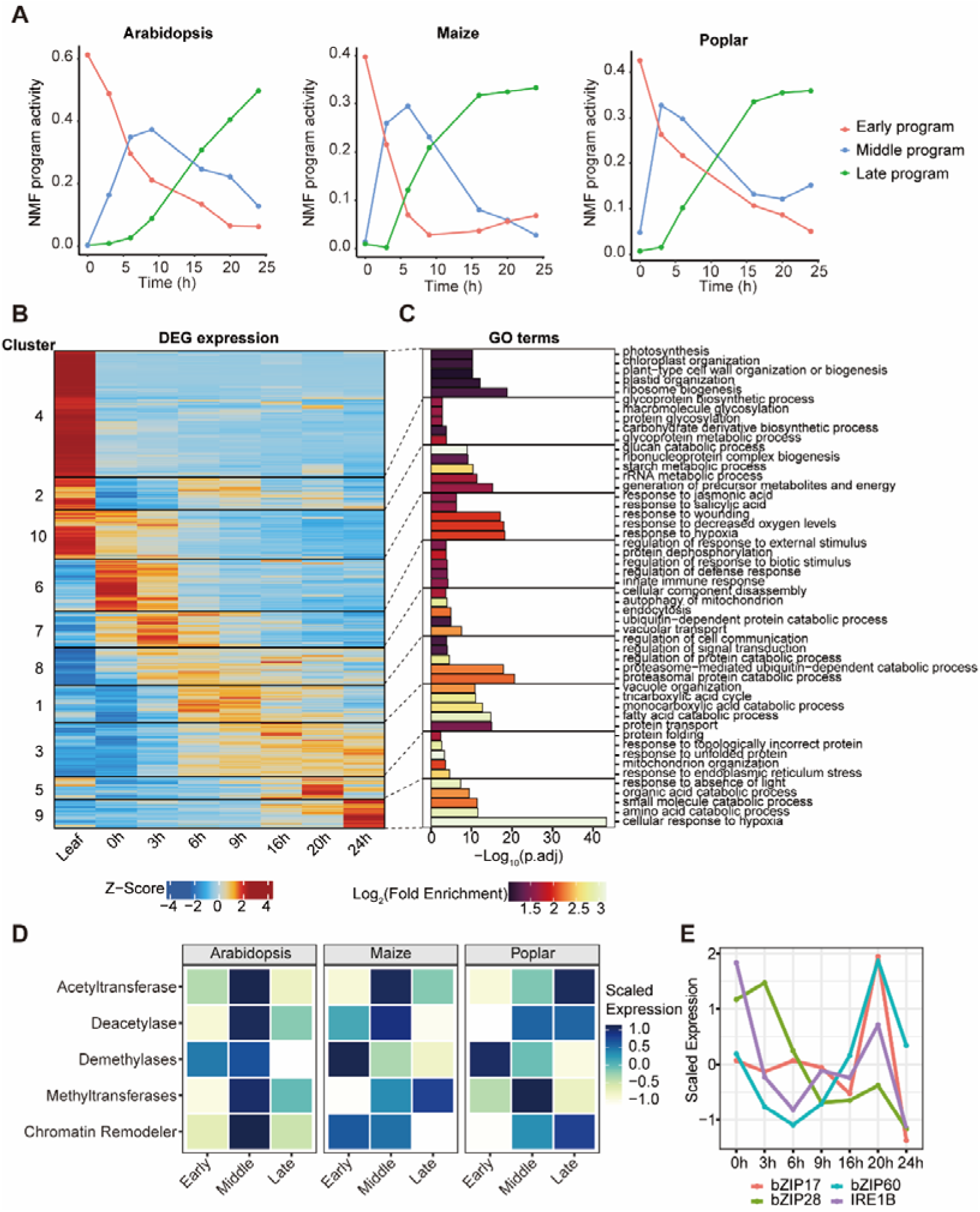
Temporal transcriptional programs during protoplast incubation. **A.** Line plots showing the activity of the three NMF-derived transcriptional programs across three species. **B.** Heatmap showing K-means clustering of DEGs in *Arabidopsis*. **C.** GO enrichment for each cluster. **D.** Heatmap showing expression of chromatin regulation related genes across three species. **E.** Line plot showing expression of ER stress gene in *Arabidopsis*.

To explore patterns of transcription in protoplast at different timepoint, we performed *k*-means clustering on the DEGs (**Supplementary Table S2**) in three species. Inspection of these clusters identified four major groups corresponding to leaf-specific genes and genes associated with early, middle, and late transcriptional responses. Leaf-specific genes were identified in *Arabidopsis* (clusters 4, 2, and 10; **Fig. 2B**), maize (clusters 8, 3, and 6; **Fig. S5A**), and poplar (clusters 1, 7, and 8; **Fig. S5C**). Early-response genes were found in *Arabidopsis* (clusters 6 and 7; **Fig. 2B**), maize (clusters 1 and 7; **Fig. S5A**), and poplar (cluster 2; **Fig. S5C**). Middle-response genes included *Arabidopsis* clusters 1 and 8 (**Fig. 2B**), maize clusters 9 and 2 (**Fig. S5A**), and poplar cluster 4 (**Fig. S5C**). Late-response genes comprised *Arabidopsis* clusters 3, 5, and 9 (**Fig. 2B**), maize cluster 4 (**Fig. S5A**), and poplar clusters 3 and 6 (**Fig. S5C**). We then performed GO enrichment analysis for each cluster within each species to characterize their functional properties. Leaf-associated clusters were enriched in photosynthesis and chloroplast organization. Early-response clusters were associated with stress responses, whereas middle-response clusters were enriched in metabolic processes in *Arabidopsis* and maize, and in hormone-related processes in poplar. Late-response clusters were enriched in stress-related processes, metabolic pathways, and protein degradation pathways (**Fig. 2C**, **Fig. S5B** and **Fig. S5D**).

As the transcriptome changed substantially, we focused on genes involved in chromatin regulation, including histone acetyltransferases, deacetylases, methyltransferases, demethylases, and chromatin remodelers (Cheng *et al*., 2020; Huang *et al*., 2025; Pandey *et al*., 2002). We found that chromatin-modifying enzymes/factors showed stage-specific expression patterns, with higher expression at the middle stage in *Arabidopsis* and maize, and a broader middle-to-late stage expression in poplar (**Fig. 2D, Supplementary Table S3**). At the family level, histone acetyltransferases and deacetylases were predominantly expressed at the middle stage in *Arabidopsis* and maize, whereas in poplar deacetylases showed middle-stage enrichment and histone acetyltransferases peaked at the late stage (**Fig. S6A-C**). Chromatin remodelers displayed a similar pattern, with higher expression at the middle stage in *Arabidopsis* and maize, but no clear early-stage enrichment in poplar (**Fig. S6-C**). In contrast, histone methyltransferases displayed a peak across species in 3–6 h (**Fig. S6A-C**). In addition, ER stress-related genes showed distinct temporal activation patterns among species. In *Arabidopsis*, they were activated at both early and late stages (**Fig. 2E**), whereas in maize they were primarily induced at the early stage. In poplar, different subsets of genes were activated at either the early or late stage (**Fig. S6D**, **E**). These results suggest coordinated but temporally distinct regulation of chromatin modification and ER stress responses across species.

### Species-specific dynamics of cell states during protoplast incubation

Given these dynamic transcriptional changes, we next asked how cellular states are altered during protoplast incubation. We first identified DEGs in different cell type from publicly available single-cell RNA datasets (Giabardo *et al*., 2026; Procko *et al*., 2022; Sun *et al*., 2022) and refined them to define robust cell-type-specific programs for each cell type in leaf protoplasts across three species (**Supplementary Table S4**). Next, we calculated the deviation (root mean square error, RMSE) of each cell type relative to the 0 h reference across three species and summarized these values to characterize changes across distinct cell-type-specific categories. In *Arabidopsis* and maize, the global deviations increased rapidly and reached a plateau within 6 h, whereas in poplar they exhibited a more gradual temporal progression (**Fig. 3A**). Consistent with this pattern, correlation analysis of cell-type-specific programs showed that samples from 6 to 24 h clustered together in *Arabidopsis* and maize, while changes in poplar were more gradual (**Fig. S7A**). In addition, different cell types exhibited distinct temporal trends in deviation dynamics. In *Arabidopsis*, phloem parenchyma cells showed more rapid transcriptional deviation, exhibiting large deviations at 3 h, whereas bundle sheath cells exhibited minimal changes (**Fig. 3B**). In maize, cell-type-specific programs exhibited rapid and substantial transcriptional changes across all cell types (**Fig. S7B**). In poplar, trichome-specific programs remained relatively stable across different time points (**Fig. S7C**). Moreover, many genes in cell-type-specific programs displayed significant transcriptional changes relative to the 0 h reference. In *Arabidopsis*, senescing cell program genes were upregulated from 6 h. In maize, a large number of differentially expressed genes were observed in epidermal and vascular cell programs, whereas in poplar, mesophyll-associated programs showed extensive transcriptional changes (**Fig. S7D**). These results suggest that tissue context is required to maintain stable cell-type-specific programs in plant cells.

**Figure 3.**
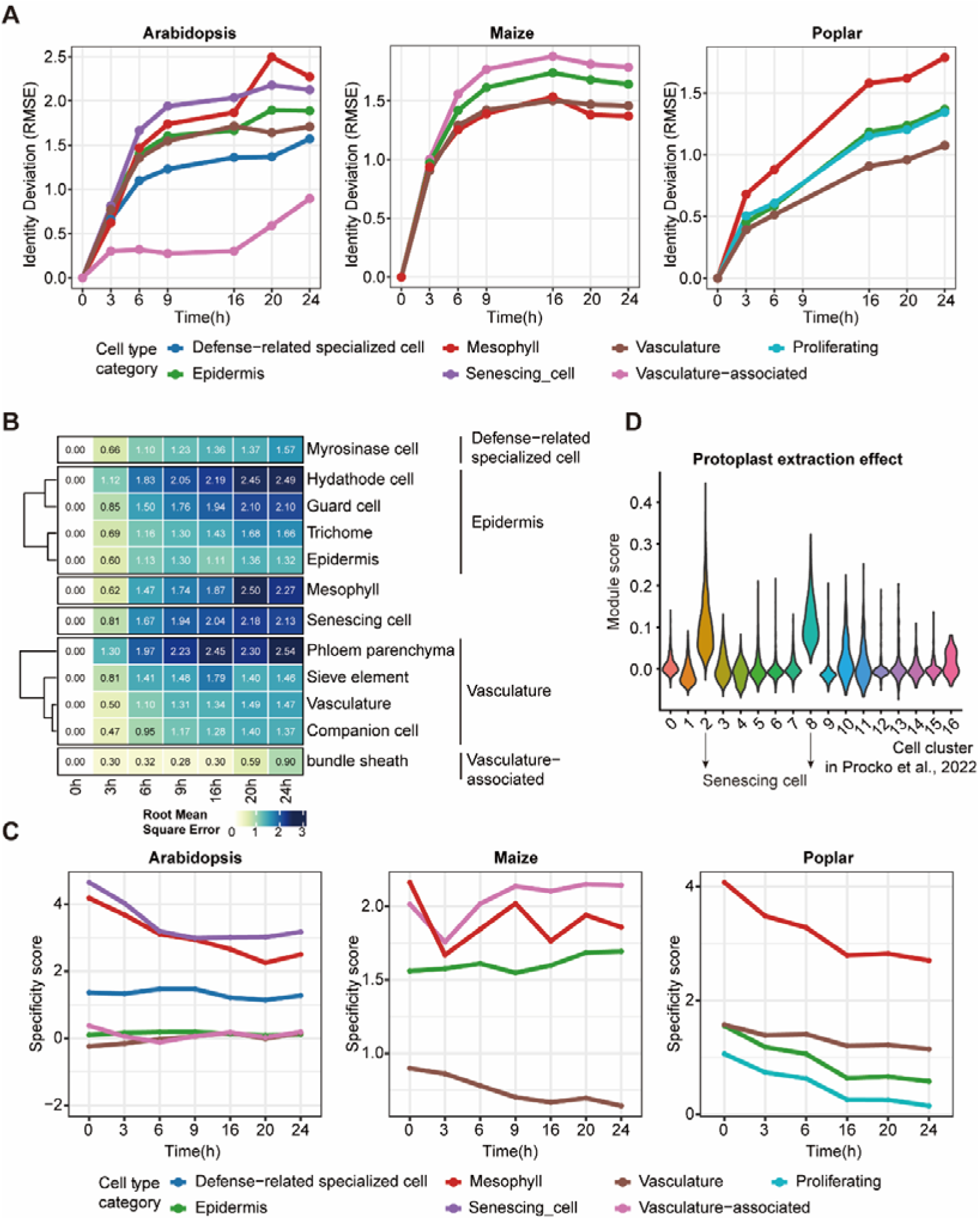
Cell-type-specific programs transition during protoplast incubation. **A.** Temporal dynamics of cell category programs deviation relative to 0h across three species. **B.** Heatmap showing cell-type-specific programs deviation for each cell types in *Arabidopsis*. **C.** Line plot showing specificity of cell-type-specific programs over time across three species. **D.** The module score of protoplast extraction effect in different cell clusters from (Procko *et al*., 2022).

Additionally, we defined a specificity score to quantify the extent to which cell-type-specific programs were enriched in each cell type relative to background levels, and tracked the temporal dynamics of this score across species (**Fig. 3C**). In *Arabidopsis*, mesophyll and senescing cell programs showed a marked decrease in specificity scores over time, whereas epidermis, vasculature-associated, defense-related, and proliferating cell programs were relatively stable or showed only mild declines. In maize, mesophyll programs displayed temporal fluctuations but ultimately decreased, whereas vasculature programs showed a gradual decline. In contrast, poplar exhibited a global decline in cell-type-specific programs, with most programs showing continuous decreases in specificity scores. These results suggest that distinct cell-type-specific programs exhibit differential sensitivity to temporal changes, with heterogeneous dynamics in transcriptional specificity across cell types and species.

### Integrative analysis reveals stress-associated cell states in protoplast-derived scRNA-seq data

Given that some single-cell data were derived from protoplasts, we leveraged bulk RNA-seq data to characterize the effects of protoplast isolation and to facilitate the identification of potential low-quality cell populations. For downstream analyses, we focused on *Arabidopsis* and used the scRNA-seq dataset (Procko *et al*., 2022) from leaf protoplasts as a reference. To examine the effects of protoplast extraction, we assessed gene activity in the reference scRNA-seq dataset, focusing on genes specifically upregulated in fresh protoplasts (cluster 6 in **Fig. 2B**). We found that these genes exhibited the highest activity in clusters 2 and 8, which were annotated as senescing cells (**Fig. 3D**; **Fig. S8A**). These findings suggest that clusters 2 and 8 capture senescence- or stress-associated states likely induced by protoplast isolation.

As most cell-type-specific programs were already largely changed at 6 h in protoplasts, we hypothesized that stressed cells may preferentially activate transcriptional programs characteristic of the 6 h and later stages. To test this, we leveraged the DEGs identified above and assessed the activity of middle (clusters 1 and 8 in **Fig. 2B**) and late (clusters 3, 5, and 9 in **Fig. 2B**) programs across different cell clusters. We next applied AUCell and Seurat’s AddModuleScore to quantify gene program activity at the single-cell level. Both methods consistently showed that clusters 14 and 15, which represent hydathode and myrosinase cells, exhibited higher activity of the middle-stage and late-stage (**Fig. S8B-D**) programs, suggesting these clusters are enriched for middle- and late-stage protoplast transcriptional programs. Together, these results demonstrate that this framework can help identify stress-associated cell states and disentangle protoplast-induced transcriptional programs in scRNA-seq data.

### Attenuation of cell-type-specific transcriptional structure at 6 h

In light of the substantial changes in cell-type-specific programs observed at 6 h, we isolated nuclei from maize leaf protoplasts at this time point and performed snRNA-seq. Quality control metrics showed low mitochondrial- and chloroplast-derived transcript fractions across all clusters (**Fig. S9A; Supplementary Table S5**), along with good UMI and gene detection rates. A strong positive correlation between UMI counts and detected genes (**Fig. S9B**) further supports consistent library complexity and overall data quality. The dataset included 3,765 cells, with a median of 1,964 unique transcripts and 1,173 genes detected per cell. By uniform manifold approximation and projection (UMAP) for dimensionality reduction, we identified 5 closely related clusters (**Fig. 4A**).

**Figure 4.**
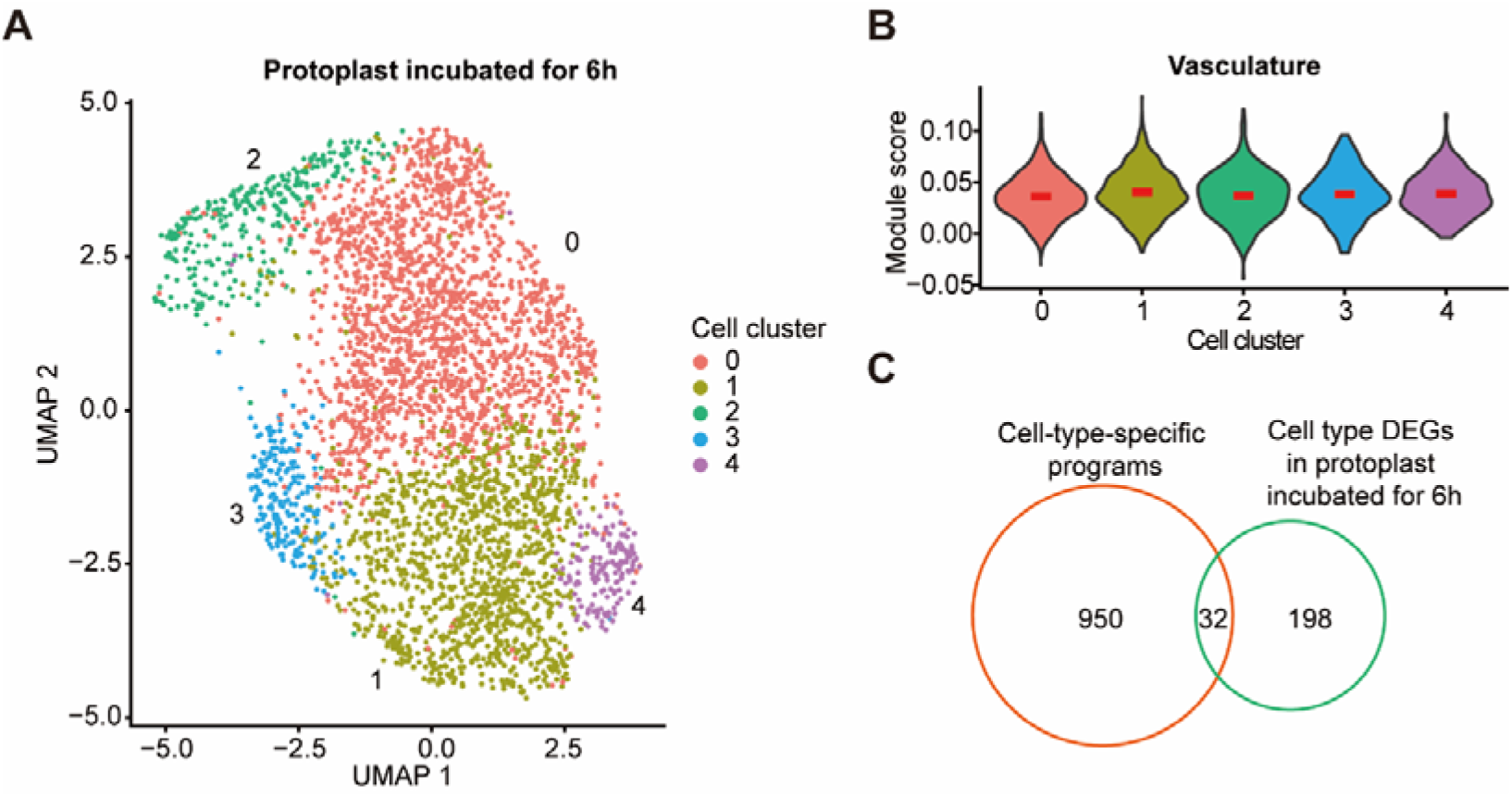
Cell-type-specific transcriptional structure attenuation in 6 h snRNA-seq. **A.** UMAP clustering of single-nucleus RNA-seq data from 6h maize protoplasts. **B.** The module score of vasculature-related genes in different cell clusters. **C.** The overlap between cell-type-associated programs and differentially expressed genes in 6 h protoplasts.

To examine cell state dynamics at 6 h, module scores were calculated for previously defined cell-type-specific programs using Seurat’s AddModuleScore function. We did not observe clear cluster-specific patterns, as cell-type-specific program activities were broadly similar across clusters (**Fig. 4B**; **Fig. S9C-D**), suggesting a reduction in cell-type-specific programs compared to fresh protoplasts. Consistently, differential expression analysis for each cluster identified relatively few DEGs, with minimal overlap with known cell-type-specific programs (32 of 982) (**Fig. 4C**). These results revealed a widespread convergence of cell-type-specific program activities at 6 h, suggesting a substantial change in cellular states.

## Discussion

We constructed a time-resolved transcriptomic dataset of protoplasts sampled at multiple time points following isolation. Our cross-species analysis reveals that transcriptomes of protoplasts are shaped by two major forces: a rapid and conserved isolation-induced transcriptional reprogramming and a progressive, time-dependent transcriptional trajectory, characterized by a rapid convergence of cell-type-specific programs within 6 hours. It also provides a framework for identifying and filtering isolation stress–associated cells in single-cell RNA-seq datasets. Together, these results establish a generalizable framework for understanding the transcriptional consequences of protoplast isolation and for improving the accuracy of single-cell RNA-seq analyses by minimizing artifacts introduced during sample preparation.

Plant cells exhibit substantial transcriptional and chromatin-level alterations during generation of protoplasts. Beyond the initial isolation response, we identified early, middle, and late transcriptional programs shared across the three species, reflecting widespread gene expression changes over time. Chromatin regulation–related genes were upregulated during the middle stage in *Arabidopsis*, maize, and poplar, and remained elevated into the late stage in poplar. Despite species-specific differences in timing and duration, their consistent induction suggests dynamic chromatin change during cell incubation. This is consistent with previous findings that protoplast isolation induces transcriptomic chaos, accompanied by a genome-wide increase in chromatin accessibility (Xu *et al*., 2021a). Importantly, protoplasts have also been shown to re-enter the cell cycle and regenerate whole plants under defined conditions, accompanied by extensive transcriptional reprogramming and chromatin remodeling, including changes in histone variant composition and epigenetic states (Chupeau *et al*., 2013). Therefore, based on our findings and previous studies, the stress-induced transcriptional and chromatin dynamics triggered by cell isolation may represent not only an acute stress response but also an early permissive state for dedifferentiation (Gomez-Cano *et al*., 2026), depending on culture conditions.

Unlike animal cells, plant cell identity is strongly dependent on positional information rather than lineage (Scheres, 2001; van den Berg *et al*., 1995; Waites and Simon, 2000; Xu *et al*., 2021b; Yu *et al*., 2017). This suggests that removal from the native tissue environment may profoundly alter transcriptional states and developmental competence, leading to destabilization or reconfiguration of established cell identity programs (Efroni *et al*., 2016; Fehér, 2015). Our data showed cell-type-specific programs were substantially altered within 6 hours in *Arabidopsis* and maize, whereas poplar exhibited a more gradual dynamic. Single-nucleus RNA-seq further confirmed that this alteration is accompanied by convergence of cell-type-specific programs. Collectively, these findings demonstrate that protoplast isolation is not a neutral sampling procedure, but rather induces a progressive loss of native cellular states during incubation, thereby diminishing the resolution of canonical cell type annotation. These observations also provide a rationale for the use of transcriptional inhibitors during protoplast preparation for plant single-cell transcriptomic studies (Tenorio Berrío *et al*., 2025; Wu *et al*., 2024), as such approaches may help preserve native transcriptional states and reduce dissociation-induced transcriptional reprogramming. More broadly, our results underscore the critical role of tissue context in maintaining plant cellular states and provide insights into how differentiated states are preserved or reshaped following cellular dissociation and the loss of tissue context.

In addition, by integrating bulk RNA-seq, reference scRNA-seq datasets, and gene program scoring, we identified distinct stress-associated cell populations that emerge during protoplast preparation and incubation. These cells exhibit activation of middle- and late-stage transcriptional programs and are enriched in senescence- or stress-like states, suggesting that they represent aberrant cell states induced by the experimental procedure. Together, these results provide a systematic framework to disentangle true biological cell states from protoplast-induced transcriptional artifacts.

Cell-based assays in plants, particularly when coupled with single-cell genomics, offer a powerful framework for dissecting gene function and cellular heterogeneity at high resolution (Zhang and Schmitz, 2025; Zhang *et al*., 2026). However, our results indicate that protoplast isolation and post-isolation incubation induces rapid and widespread transcriptional dynamics, including stress responses and convergence of cell-type–specific programs, which may complicate direct interpretation of native *in vivo* states. These findings highlight the importance of rapid sample processing in protoplast-based systems and emphasize the need for careful temporal consideration when inferring endogenous cellular programs, while also providing a framework for interpreting plant cellular state maintenance and identifying stress-associated transcriptional states in protoplast-based systems.

## Methods

### Plant Growth in *Arabidopsis* and Maize

*Arabidopsis thaliana* plants (Columbia-0, Col-0) were sown in soil and grown in a growth chamber at 22 °C under a 16 h light/8 h dark photoperiod, after 4 weeks growth, and well-expanded leaves were collected for protoplast isolation. Maize B73 seeds were sown in soil and grown in a growth chamber at 25–28 °C under a 16 h light/8 h dark photoperiod. Once the coleoptiles emerged from the soil surface, the trays were transferred to a dark environment at room temperature. After 10 days of growth from sowing, maize leaves were collected for protoplast isolation.

### Poplar tissue culture and maintenance

Poplar nodal explants (P. tremula × P. alba, INRA 717-1B4) were cultured on shoot elongation medium (SEM) to induce axillary shoot growth. The SEM consisted of basal Murashige and Skoogs (MS) medium supplemented with 3% sucrose, 0.2 g/L glutamine, and 0.05 mg/L 6-Benzylaminopurine (BAP), pH 5.8. Nodes were placed on SEM and maintained under standard at 25°C with 16h of light until new shoots emerged from the nodal explants. After approximately 2–3 weeks, regenerated shoots were transferred to root induction medium (RIM). The RIM consisted of half-strength MS medium supplemented with 2% sucrose, 0.2 g/L glutamine, and 0.1 mg/L Indole-3-butyric acid (IBA), pH 5.8. Shoots were maintained on RIM for 3-4 weeks to promote rooting and plantlet development. For protoplast isolation, young well expanded leaves were collected from healthy in vitro-grown plants after 3-4 weeks on RIM. Leaves were harvested from actively growing plants and used immediately for downstream protoplast isolation.

### Protoplast isolation in *Arabidopsis*

Following the protocol described by (Yoo *et al*., 2007), the middle part of leaf was cut into small strips using a sharp razor blade, then leaf strips were transferred into the prepared enzyme solution containing 20 mM MES (pH 5.7), 1.5% (w/v) cellulase R10, 0.4% (w/v) macerozyme R10, 0.4 M mannitol, 20 mM KCl, 10 mM CaCl□, 2 mM β-mercaptoethanol, and 0.1% bovine serum albumin (BSA). The leaf strips were vacuum infiltrated for 30 min in the dark, with intermittent release to enhance infiltration and then incubated without shaking in the dark at room temperature for at least 3 h. An equal volume of W5 solution (2 mM MES, pH 5.7, containing 154 mM NaCl, 125 mM CaCl□, and 5 mM KCl) was added to dilute the enzyme/protoplast mixture, which was then filtered through a 75-μm nylon mesh. The filtrate was centrifuged at 100 × g at 4 °C for 2 min to pellet the protoplasts. After removal of the supernatant, the protoplasts were resuspended in W5 solution for washing and centrifuged again at 100 × g at 4 °C for 2 min to pellet the cells. The supernatant was then discarded, and the protoplasts were resuspended in WI solution (4 mM MES, pH 5.7, containing 0.5 M mannitol and 20 mM KCl) and transferred to individual wells of a 6-well tissue culture plate covered with aluminum foil, then incubated at room temperature.

### Protoplast isolation in maize

The middle portion of the second leaf from maize seedling was selected and cut into strips, which were then transferred into enzyme solution containing 10 mM MES (pH 5.7), 2% (w/v) cellulase R10, 0.6% (w/v) macerozyme R10, 0.6 M mannitol, 1 mM CaCl□, 5 mM β-mercaptoethanol, and 0.1% (w/v) BSA. The leaf strips were vacuum infiltrated for 20 min in the dark, with intermittent release to enhance infiltration and then incubated with gentle shaking (30–40 rpm) in the dark at room temperature for 2 h. After 2h rotation, the plate was rotated at 80 RPM for 5 minutes to release protoplasts and strained through 40 μm strainer to remove undigested tissues. Centrifugation was performed at 100 × g for 3 min to pellet the protoplasts. The supernatant was removed and the protoplasts were washed with 10 mL chilled Mmg buffer (0.6 M mannitol, 4 mM MES, pH 5.7, and 15 mM MgCl□). The suspension was centrifuged again under the same conditions, and the protoplasts were finally resuspended in WI solution (0.5 M mannitol, 4 mM MES, pH 5.7, and 20 mM KCl) and transferred to individual wells of a 6-well tissue culture plate covered with aluminum foil, then incubated at room temperature.

### Protoplast isolation in poplar

Fresh poplar leaves were cut into strips and incubated in enzyme solution containing 2% (w/v) cellulase, 0.30% (w/v) macerozyme, 0.10% (v/v) pectinase, 20 mM MES (pH 5.7–5.8), 0.4 M mannitol, 1 mM CaCl□, 0.2% β-mercaptoethanol, and 0.1% (w/v) BSA. Then leaf strips were vacuum infiltrated for 20 min in the dark, with intermittent release to enhance infiltration. After vacuum treatment, tissues were incubated on a shaker at 50 rpm for 1 h 45 min at room temperature, followed by gentle shaking at 80 rpm for 5 min to facilitate protoplast release. The digestion mixture was filtered sequentially through 70 μm and 40 μm nylon meshes and centrifuged at 150 × g for 3 min at 4 °C. After remove the supernatant, and protoplasts were resuspended in storage solution (0.4 M mannitol, 20 mM MES, pH 5.7–5.8, 1 mM CaCl□, and 0.2% β-mercaptoethanol) and transferred to individual wells of a 6-well tissue culture plate covered with aluminum foil, then incubated at room temperature.

### RNA extraction

Total RNA of protoplasts was extracted using the RNeasy UCP Micro Kit (QIAGEN, 73934) according to the manufacturer’s instructions. Briefly, tissue samples were homogenized in lysis buffer containing β-mercaptoethanol, and the lysates were applied to spin columns. Following on-column DNase digestion to remove genomic DNA contamination, the columns were washed sequentially with the provided buffers, and total RNA was eluted in RNase-free water. RNA concentration and purity were determined using a spectrophotometer, and RNA integrity was assessed prior to downstream applications.

### RNA-seq data processing

Raw paired-end RNA sequencing reads were processed using fastp (v0.23.4) (Chen *et al*., 2018) to remove adapter sequences and low-quality bases with automatic adapter detection for paired-end reads. Cleaned reads were aligned to the corresponding reference genome using HISAT2 (v2.2.1) (Kim *et al*., 2015) with the parameters “--min-intronlen 20 --max-intronlen 4000”. The reference assemblies included *Arabidopsis thaliana* (TAIR10), maize (B73 RefGen_v4), and poplar 717 (v5.1) (Zhou *et al*., 2023). Alignment files were converted to BAM format, filtered to retain reads with mapping quality scores ≥ 20, and sorted using SAMtools (v1.18) (Li *et al*., 2009). Gene-level read counts were generated using featureCounts in the Subread package (v2.0.6) (Liao *et al*., 2013) with the parameters “-t exon -g gene_id -p -P -B -C”. The resulting count matrix was used for downstream differential gene expression analyses.

Differential gene expression analysis was performed using DESeq2 (v1.42.0) (Love *et al*., 2014) in R. Raw gene count matrices generated by featureCounts were used as input. Genes with an adjusted P value (Benjamini–Hochberg false discovery rate) < 0.05 and an absolute log2 fold change > 1 were considered differentially expressed.

Gene Ontology (GO) enrichment analysis was performed using clusterProfiler (v4.12.6) (Yu *et al*., 2012) in R. For *A. thaliana*, GO enrichment was conducted using the enrichGO function with the org.At.tair.db database. For maize and poplar datasets, GO enrichment was performed using the enricher function in clusterProfiler with organism-specific gene set annotation files, using the same statistical framework. P-values were adjusted using the Benjamini–Hochberg (BH) method, and GO terms with adjusted p-value < 0.05 were considered significantly enriched.

### Time-course modeling analysis

A spline-based limma framework was used to identify genes exhibiting significant temporal expression dynamics. Sampling time (0, 3, 6, 9, 16, 20, and 24 h) was modeled as a continuous variable, and natural cubic splines with three degrees of freedom were applied to capture nonlinear temporal expression patterns. Differential expression analysis was performed using linear modeling in limma (v3.60.6), with empirical Bayes moderation used to stabilize variance estimates across genes. P values were adjusted for multiple testing using the BH method, and genes with adjusted P values < 0.05 were defined as significantly time-dependent.

### Non-negative matrix factorization analysis

The gene expression matrix was log-transformed, and low-information genes were filtered by retaining those with mean expression > 1 and variance above the 70th percentile of the overall gene variance distribution. To determine the optimal factorization rank (k) for non-negative matrix factorization, models were fitted for k = 2–5 using the Brunet algorithm with 10 random initializations per rank, and the optimal rank was selected based on the maximum cophenetic correlation coefficient. Final NMF decomposition was performed using the selected rank with the Brunet update method and 10 independent runs to ensure solution stability. The resulting coefficient (H) matrices were extracted, which reflect program activities. Temporal trajectories of NMF-derived gene expression programs were summarized by plotting mean program activity over time.

### Cell-type-specific program genes identification

Cell-type-specific program genes were identified from publicly available single-cell RNA-seq datasets across three species (Giabardo *et al*., 2026; Procko *et al*., 2022; Sun *et al*., 2022). Genes were defined as cell-type-specific if they showed significant upregulation in a given cell type (log2 fold change > 0.5, FDR < 0.05), be detected in at least 25% of cells within that population, and exhibit at least 15% higher detection frequency compared with all other cell types.

### Nucleus extraction and single-nucleus sequencing

Nuclei were isolated from 6 h protoplasts. The nuclei isolation buffer (NIB) consisted of 10 mM MES-KOH (pH 5.4), 10 mM KCl, 10 mM NaCl, 3 mM MgCl□, 0.25 M sucrose, 0.1 mM spermine, 0.5 mM spermidine, and 1 mM DTT, supplemented with 0.5% NP-40, 0.5% PVP-40, an EDTA-free protease inhibitor cocktail, RNase inhibitors (RiboLock, Protector RNase Inhibitor, and SUPERase Inhibitor), and a fixation additive consisting of 3% glyoxal and 0.75% acetic acid. The nuclei wash buffer (NIB-wash) contained 10 mM MES-KOH (pH 5.4), 10 mM KCl, 10 mM NaCl, 3 mM MgCl□, 0.25 M sucrose, 0.5% BSA, and RNase inhibitors.

Briefly, 200 µL NIB buffer was added to the protoplast suspension, followed by incubation with gentle rotation at 4°C for 5 min and vortexing for 30 s to release nuclei. The lysate was filtered through a 40 µm cell strainer and incubated on ice for 10 min for fixation. Nuclei were then pelleted by centrifugation at 500 × g for 5 min at 4°C, and the supernatant was removed. The pellet was resuspended in 600 µL NIB-wash buffer and filtered through a 20 µm strainer. Nuclei were further washed with 500 µL NIB-wash buffer and centrifuged again at 500 × g for 5 min at 4°C. Finally, approximately 20,000 nuclei were collected for downstream library construction.

Single-nuclei RNA-seq libraries were prepared using a nucleus-based *in situ* reverse transcription and droplet barcoding workflow adapted from the 10x Genomics Single Cell ATAC kit. Isolated nuclei were subjected to reverse transcription in a 25 µL reaction containing TruseqR2 oligo-dT primer, 5× NaCl RT buffer, PEG8000, dNTPs, Maxima H-minus reverse transcriptase, and RNase inhibitors (RiboLock, Protector RNase inhibitor, and SUPERase Inhibitor). Reactions were incubated at 25°C for 10 min followed by 37°C for 15 min. After *in situ* RT, nuclei were gently resuspended and washed with ice-cold DNB-BSA buffer and centrifuged at 500 × g for 5 min at 4°C, leaving nuclei for downstream processing. Barcoding, droplet generation and GEM incubation were performed using a 10x Chromium system following manufacturer’s instructions. Nuclei were combined with a barcoding master mix and loaded onto a Chromium Chip H for GEM formation. Following GEM breakage, libraries were purified using a combination of recovery agent extraction and magnetic bead cleanup (Dynabeads MyOne and SPRIselect). DNA was washed with 80% ethanol, eluted in Elution Solution I, and further purified through two rounds of SPRI size selection. Final libraries were amplified using Q5/NEBNext 2× PCR master mix with P5 and indexed P7 primers under cycling conditions: 98°C for 1 min; 15 cycles of 98°C for 20 s, 63°C for 20 s, and 72°C for 20 s; followed by 72°C for 1 min. Purified libraries were used for high-throughput sequencing.

### SnRNA-seq data processing

Raw paired-end single-cell sequencing data were processed to extract cell barcodes and unique molecular identifiers (UMIs) prior to alignment. Cell barcodes and unique molecular identifiers (UMIs) were extracted and processed from Read 2 using seqkit (v 2.9.0) (Shen *et al*., 2016). UMI sequences were identified based on a predefined pattern and extracted from the first 10 bp of matched reads. Cell barcode sequences were reverse-complemented to ensure correct orientation. Barcode and UMI information were then paired and merged using sequence name matching, and concatenated into a single barcode–UMI tag for each read, generating a combined FASTQ file for downstream analysis. Adapter trimming and quality filtering were performed using cutadapt (v4.9) (Martin, 2011), including removal of Nextera/TruSeq adapters, NextSeq-based quality trimming, poly-G trimming, and discarding reads shorter than 20 bp while maintaining paired-end synchronization.

Cleaned paired-end reads were aligned to the reference genome using STAR (v2.7.10b) (Dobin *et al*., 2013) in STARsolo mode. Read 1 contained cDNA sequences, and Read 2 contained 16 bp cell barcodes and 10 bp UMIs extracted from fixed positions. Alignment was performed using a two-pass strategy with GTF annotation, allowing only uniquely mapped reads. Cell barcodes were matched to a whitelist with up to one mismatch, and UMI deduplication was performed using a 1-mismatch collapsing strategy. Gene expression quantification was generated at the gene level, and low-quality barcodes were filtered using EmptyDrops-based cell calling. The resulting gene-by-cell matrix was used for downstream analysis.

Downstream analyses were performed with the R package Seurat (v.5.5.0) (Hao *et al*., 2024). Initial quality control was performed in Seurat by filtering low-quality cells based on UMI counts, gene counts, mitochondrial/chloroplast RNA content, and gene complexity. Cells were retained if they satisfied the following criteria: UMI counts above the 15th percentile, gene counts above a median-based lower bound (median ± MAD × 1), gene counts below an upper threshold, mitochondrial and plastid RNA content below 15%, and log10 genes-per-UMI > 0.85. To remove potential doublets, normalized data were processed using SCTransform with regression of mitochondrial and plastid content, followed by PCA (20 components) and UMAP embedding. Doublets were identified using DoubletFinder with expected doublet rates scaled by cell number. Cells classified as doublets were removed, and only singlet cells were retained for downstream analysis. Final filtered datasets were saved for subsequent analyses.

## Data availability

All data generated in this study have been deposited in the NCBI Sequence Read Archive under BioProject accession PRJNA1489660.

## Author contributions

H.Z. and R.J.S. conserved and designed the study and experiment. H.Z. performed protoplast isolation, RNA extraction, nuclei extraction, single-nuclei library generation, data analysis and wrote the manuscript. A.S. prepared tissue-cultured poplar material for protoplast isolation. A.G., J.C.W., J.B., S.S.C., J.P.H., K.M., B.V., and C.R.B. provided valuable feedback and contributed to discussions that improved the study.

## Supplementary information

Supplementary Table S1. Summary of RNA-seq sequencing and alignment statistics.

Supplementary Table S2. Lists of differentially expressed genes identified across the three species.

Supplementary Table S3. Expression patterns of chromatin-modifying enzymes and factors across three species.

Supplementary Table S4. Lists of cell-type-specific gene expression programs in leaf protoplasts across three species.

Supplementary Table S5. Summary of snRNA-seq sequencing and alignment statistics.

**Figure S1.**
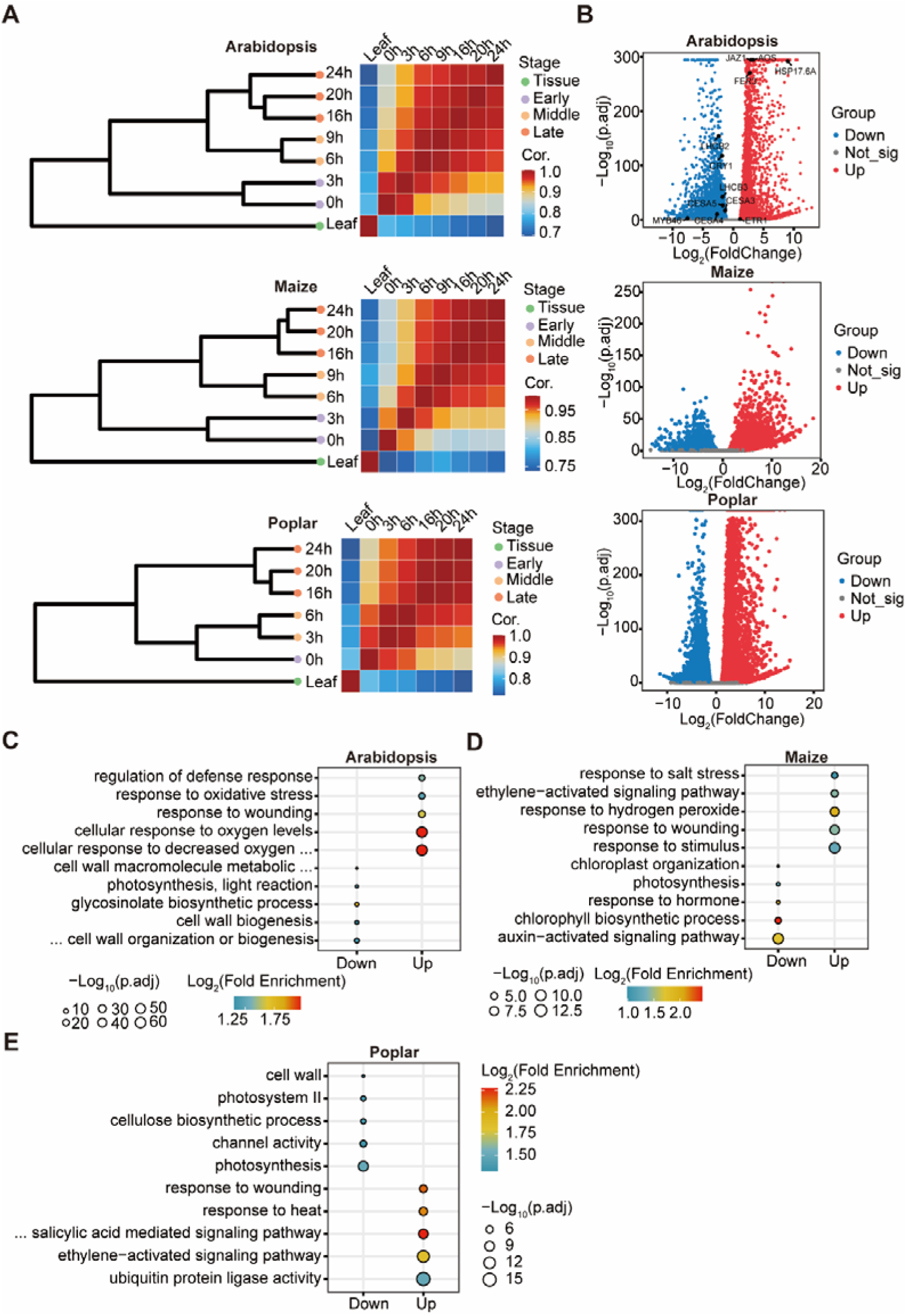
Overview of transcriptomic profiles and effects of protoplast isolation. **A.** Temporal clustering of transcriptomic profiles during protoplast incubation across three species. **B.** Volcano plot of fresh protoplast (0h) vs leaf tissue across three species. **C.** GO enrichment of DEGs induced by protoplast isolation in *Arabidopsis*. **D.** GO enrichment of DEGs induced by protoplast isolation in maize. **E.** GO enrichment of DEGs induced by protoplast isolation in poplar.

**Figure S2.**
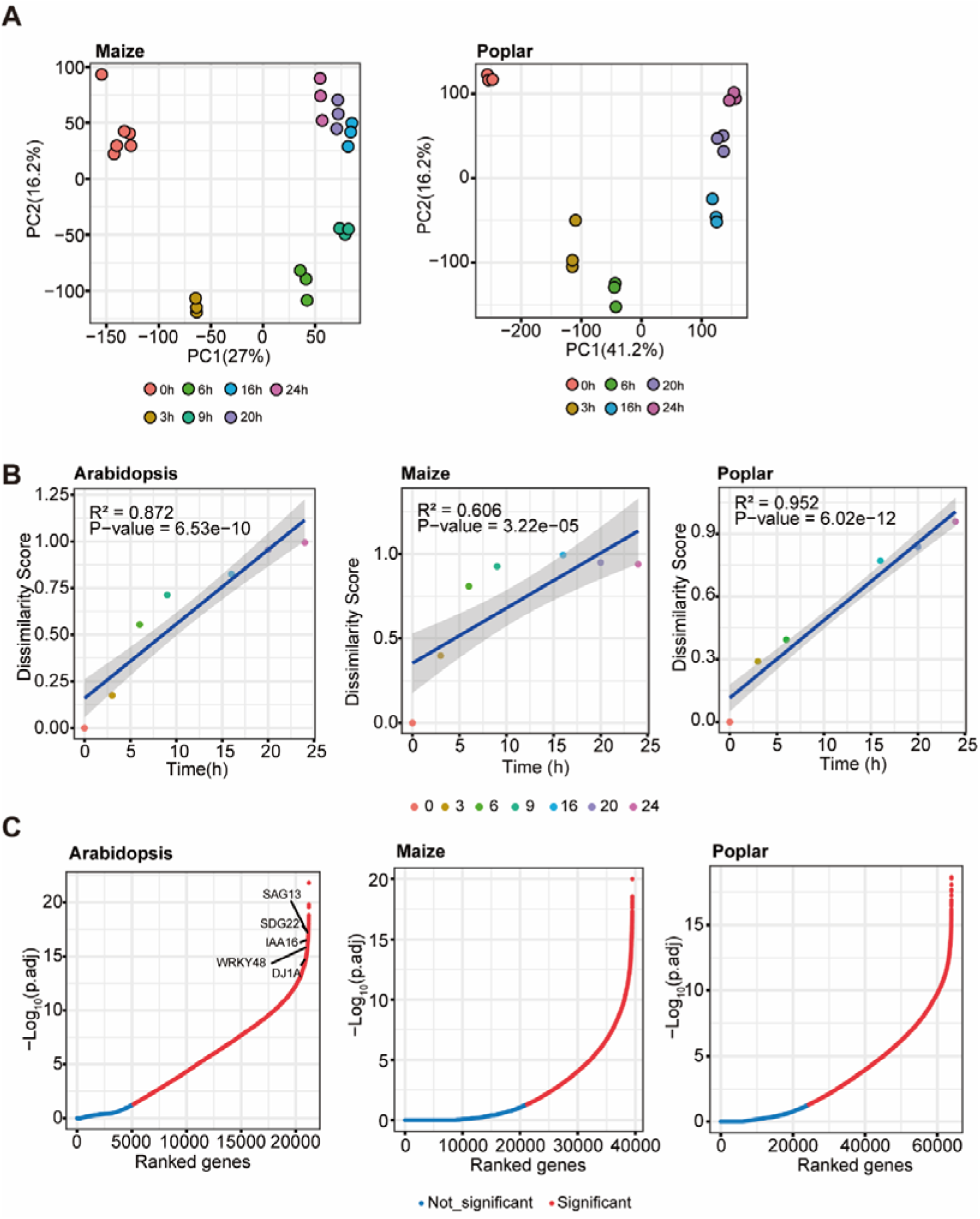
Transcription is regulated in a time-dependent manner during protoplast incubation. **A.** PCA of gene expression at different time points in maize and poplar. **B.** Linear regression of dissimilarity score against time across three species. **C.** Significance ranking of global gene expression changes in three plant species.

**Figure S3.**
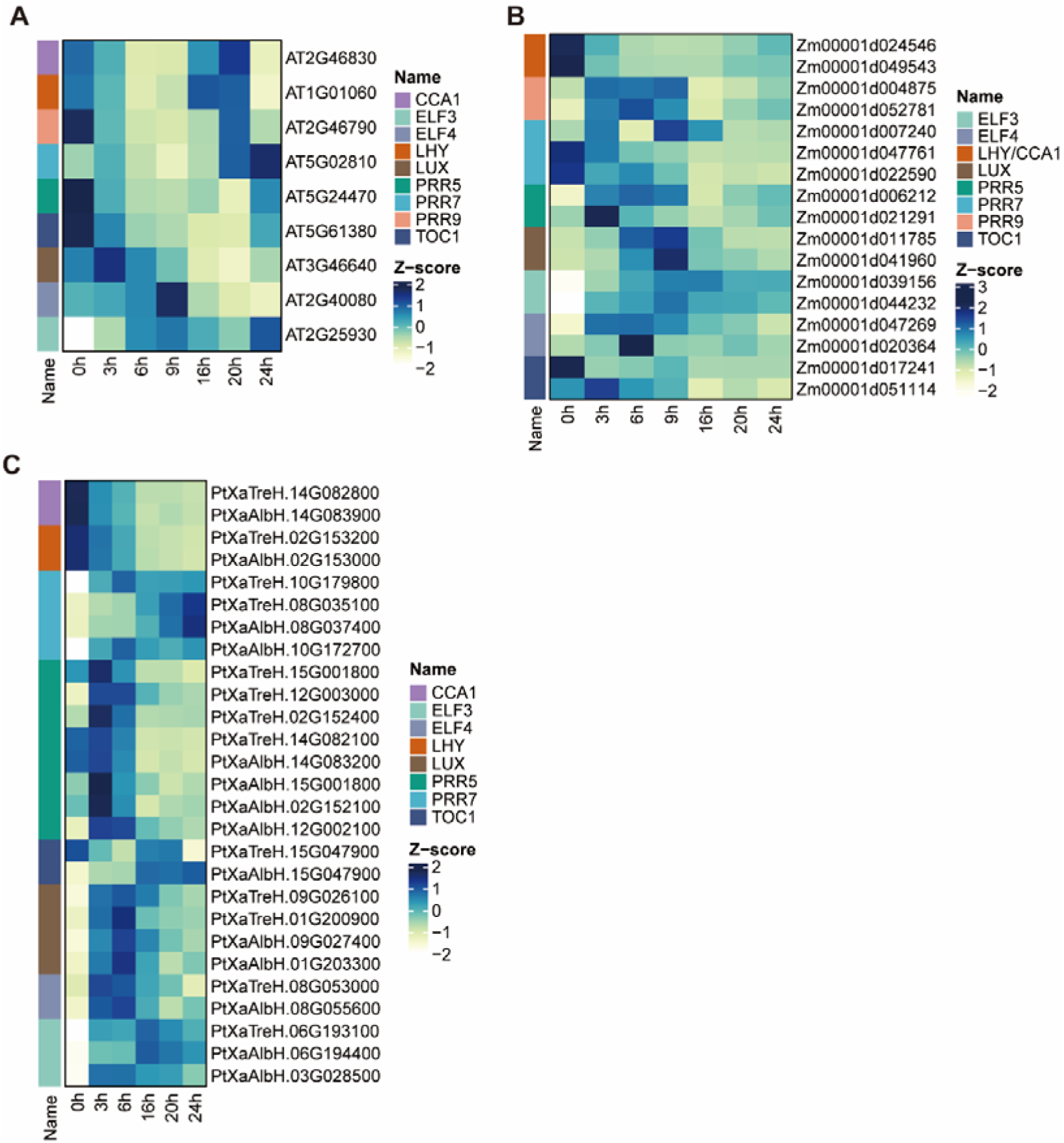
Expression patterns of circadian genes across three species. **A.** Heatmap showing expression dynamics of circadian related genes in *Arabidopsis*. **B.** Heatmap showing expression dynamics of circadian related genes in maize. **C.** Heatmap showing expression dynamics of circadian related genes in poplar.

**Figure S4.**
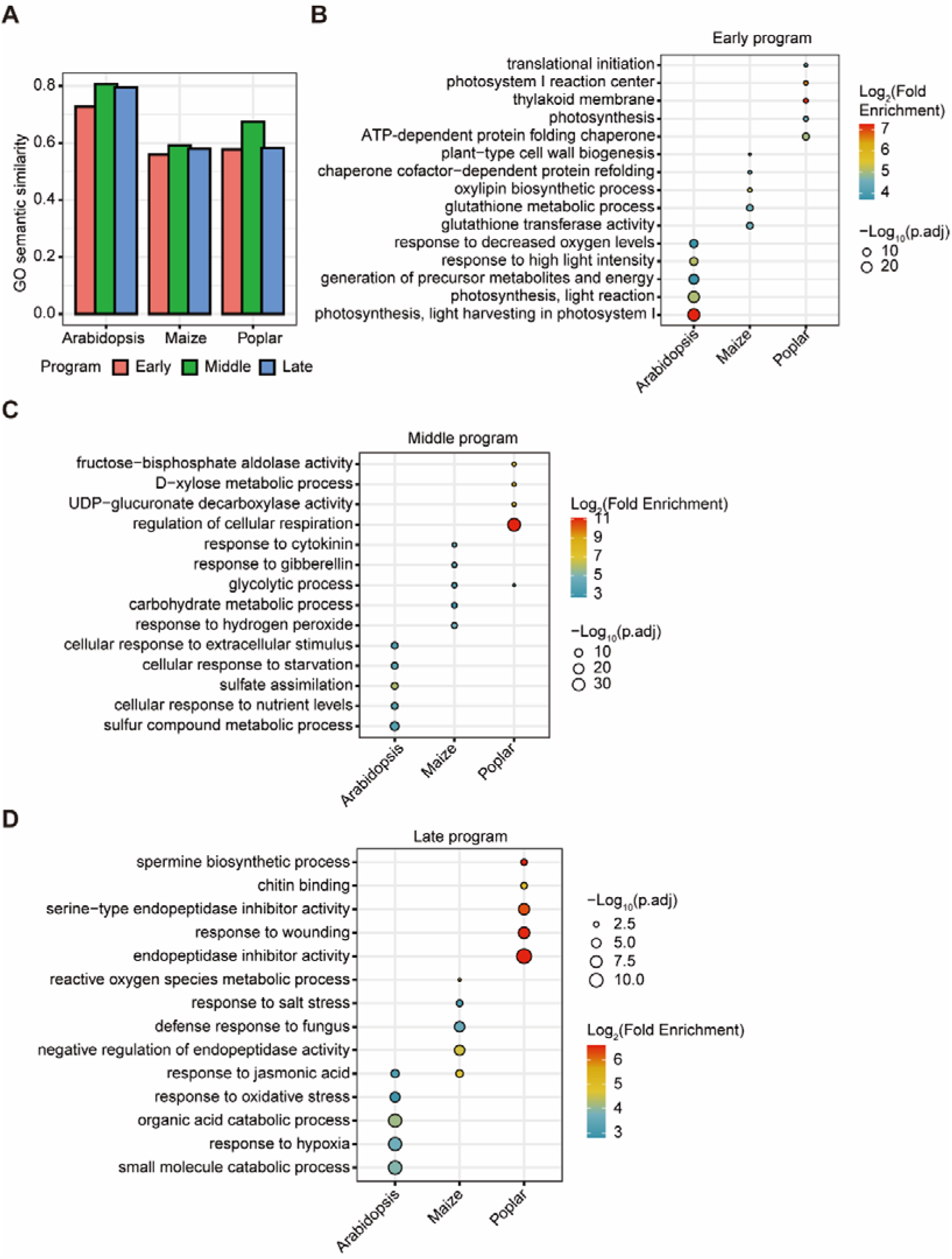
GO enrichment of early, middle, and late program across three species. **A.** GO semantic similarity between top 100 and 500 gene sets across three species **B.** GO enrichment of early program across three species. **C.** GO enrichment of middle program across three species. **D.** GO enrichment of late program across three species.

**Figure S5.**
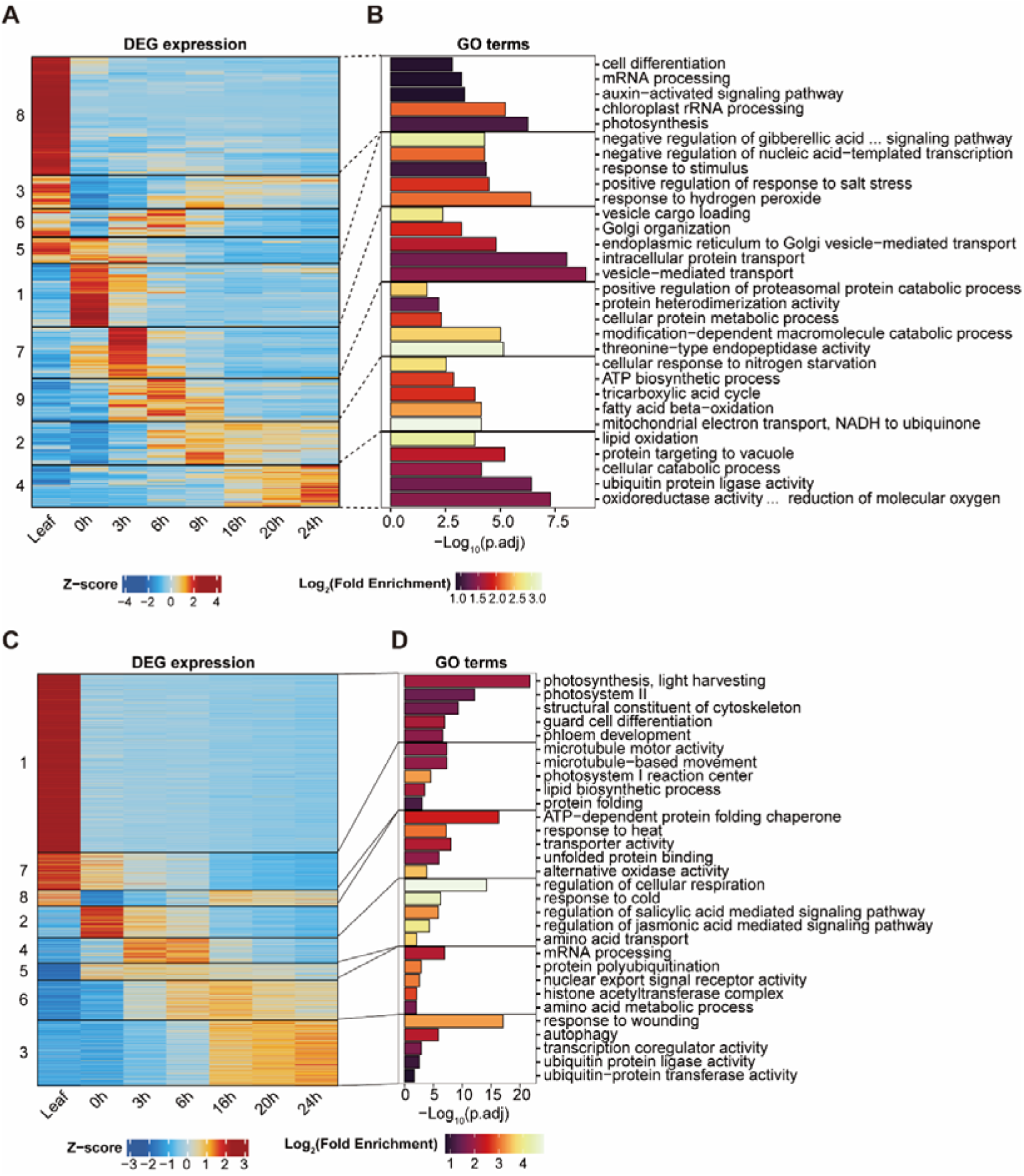
DEGs identified at different time point during protoplast incubation. **A.** Heatmap showing K-means clustering of DEGs in maize. **B.** GO enrichment for each cluster in (**A**). **C.** Heatmap showing K-means clustering of DEGs in poplar. D. GO enrichment for each cluster in (**C**).

**Figure S6.**
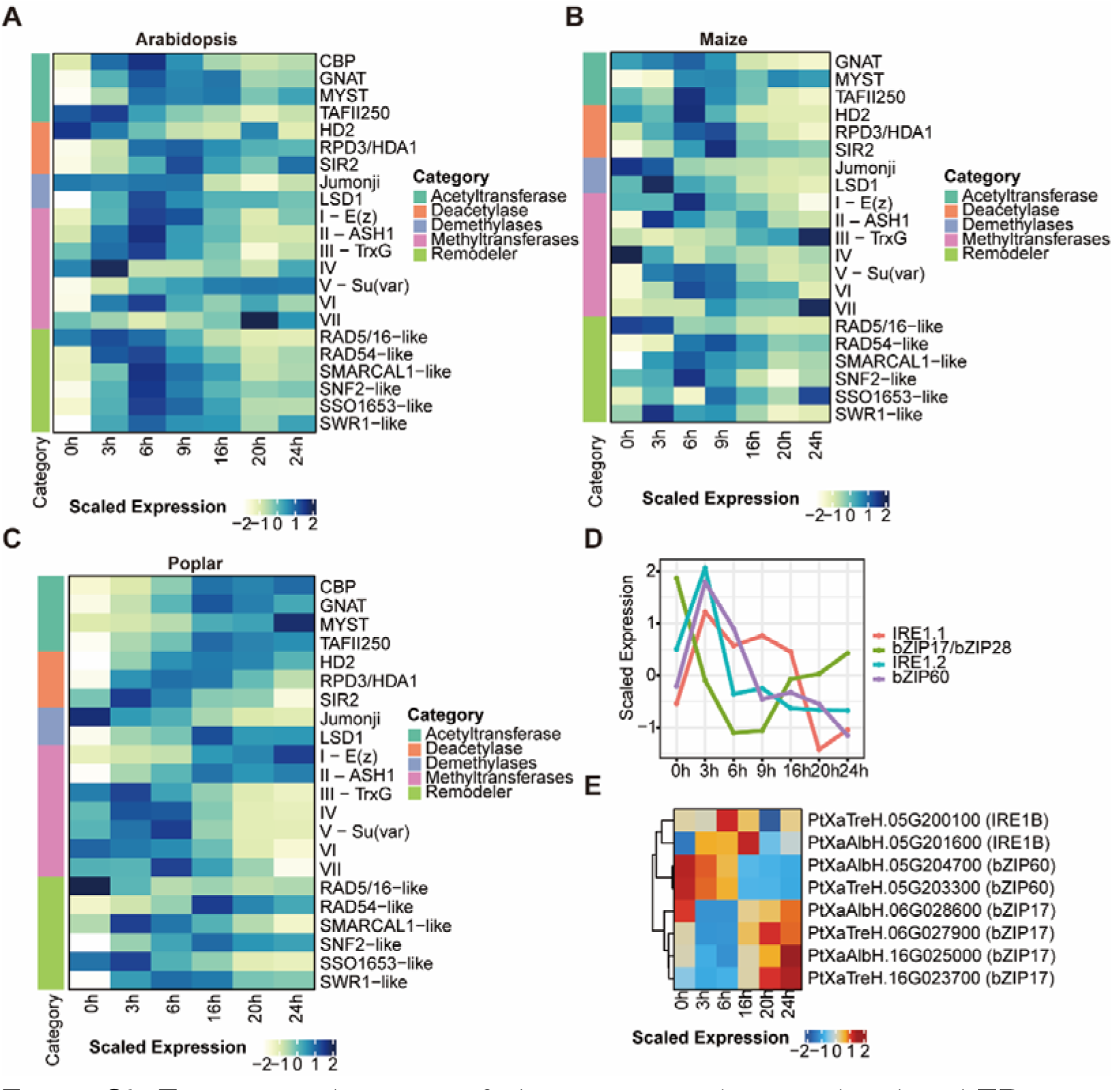
Expression dynamics of chromatin regulation related and ER stress genes across three species. **A.** Heatmap showing expression dynamics of chromatin regulation related genes in *Arabidopsis*. **B.** Heatmap showing expression dynamics of chromatin regulation related genes in maize. **C.** Heatmap showing expression dynamics of chromatin regulation related genes in poplar. **D.** Line plot showing expression dynamics of ER stress genes in maize. E. Heatmap showing expression dynamics of ER stress genes in poplar.

**Figure S7.**
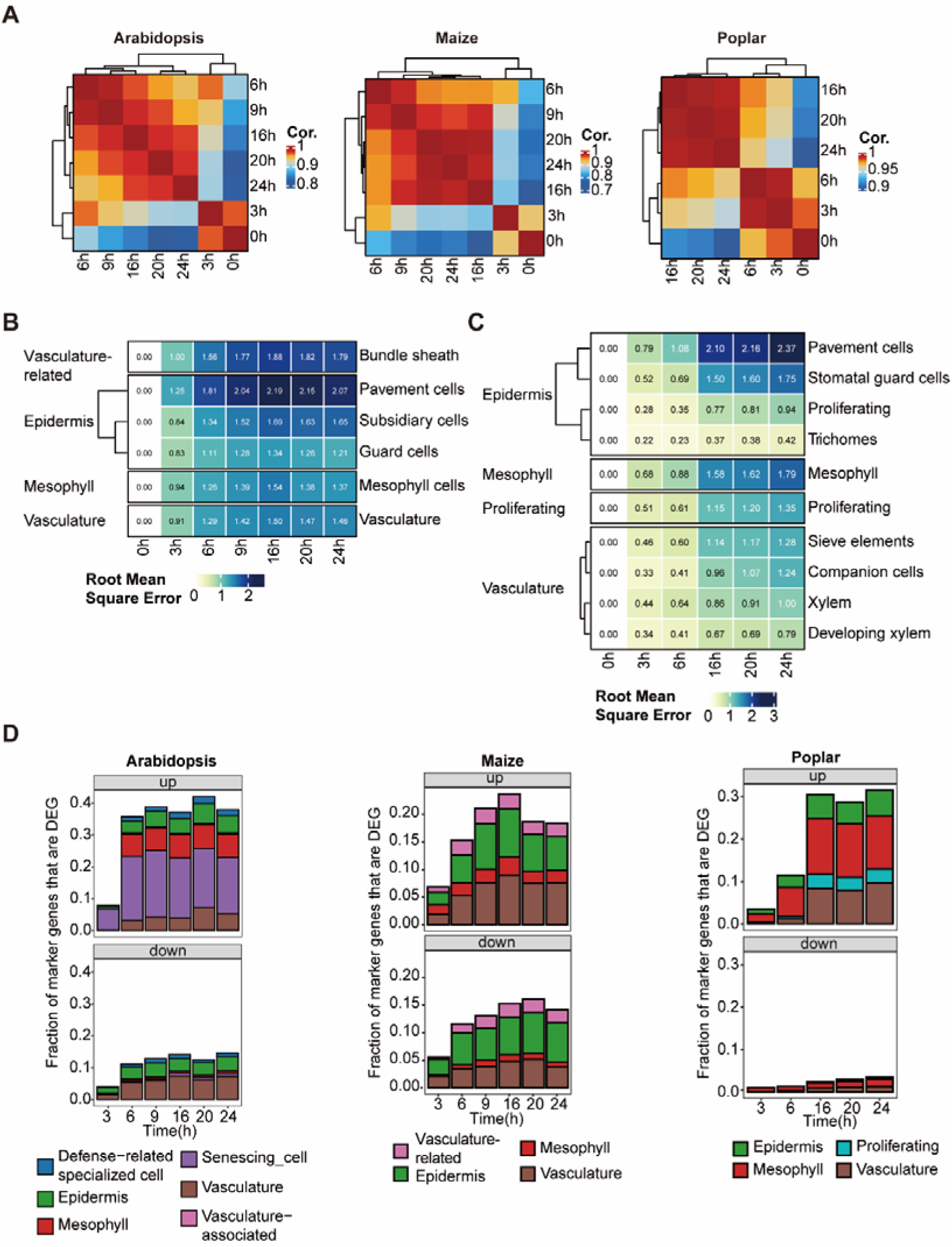
Dynamic change of cell-type-specific programs during protoplast incubation. **A.** Correlation heatmap of cell-type-specific genes across time points in three species. **B.** Heatmap showing deviations of cell-type-specific programs in maize. **C.** Heatmap showing deviations of cell-type-specific programs in poplar. **D.** Proportion of cell-type-specific genes among DEGs compared to 0h across the three species.

**Figure S8.**
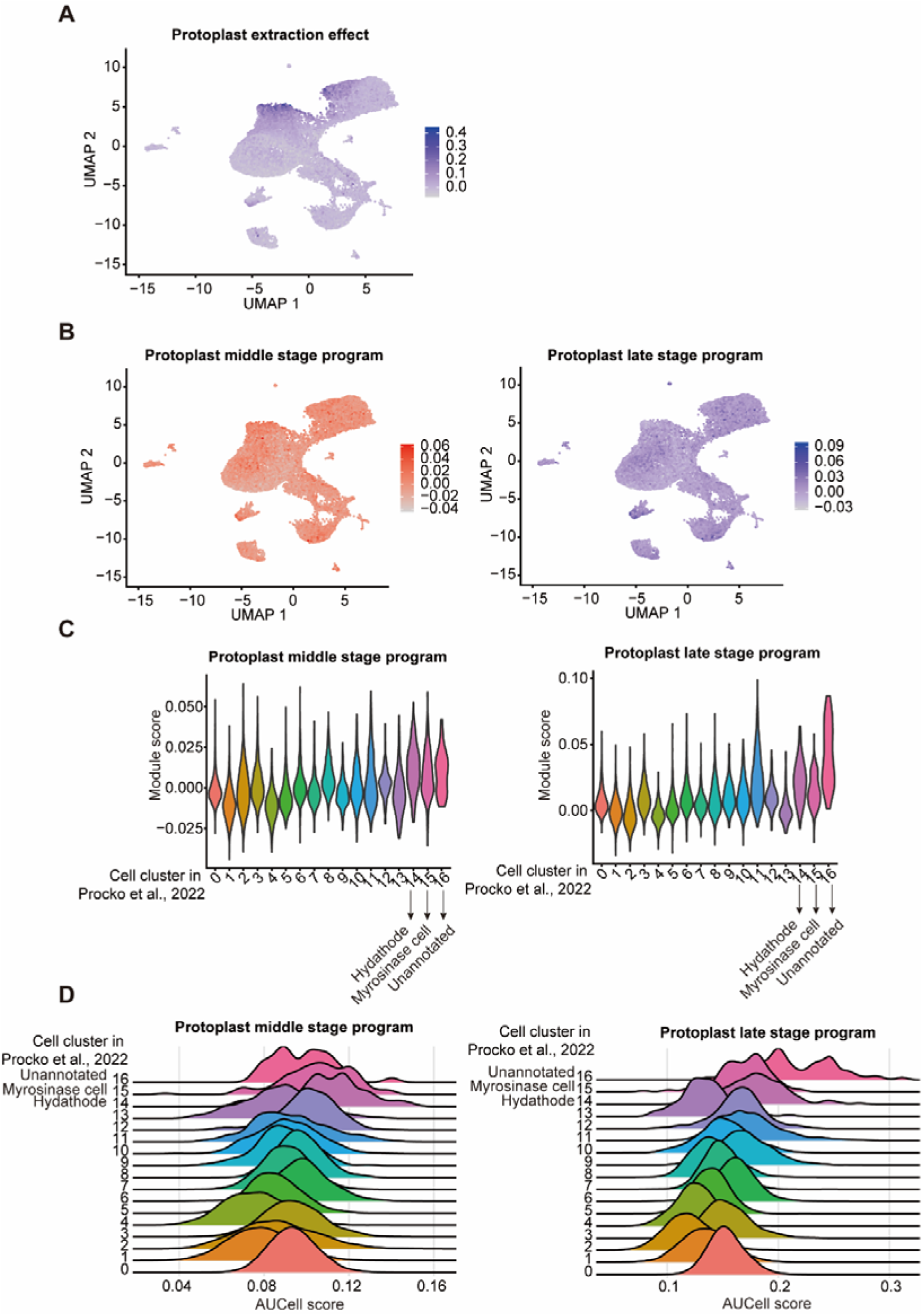
Integration of bulk RNA-seq data with published scRNA-seq data (Procko *et al*., 2022). **A.** Average expression of protoplast extraction related genes in the published scRNA-seq dataset. **B.** Module scores for middle and late stage program related genes projected on UMAP embedding. **C.** The module score of protoplast middle and late stage program in different cell clusters from published scRNA-seq data. **D.** Distribution of AUCell scores for the “middle-stage” program across cell clusters in (Procko *et al*., 2022).

**Figure S9.**
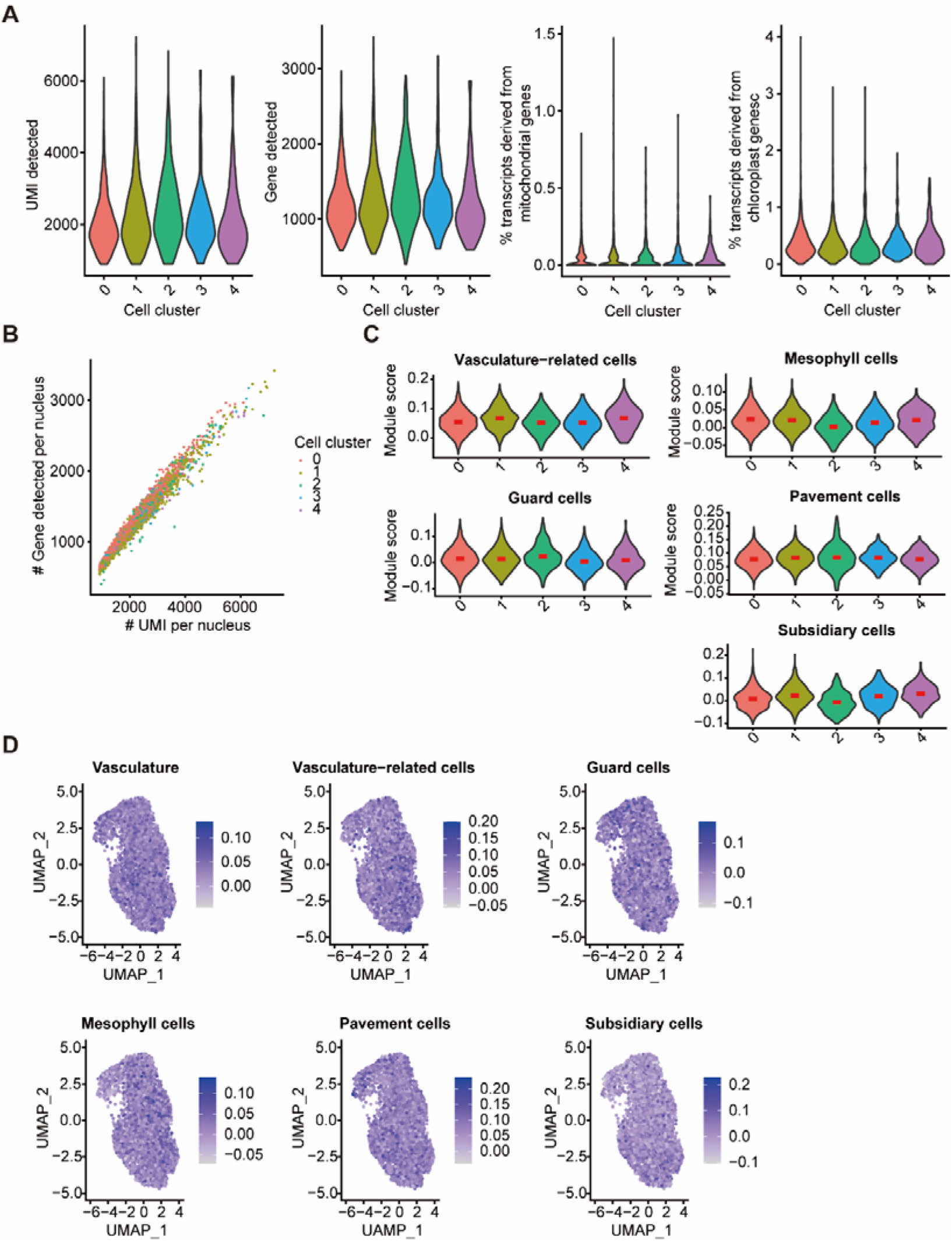
Overview of single-nuclei datasets from 6 h maize protoplasts. **A.** Quality control metrics for snRNA-seq dataset. **B.** Transcriptional complexity of nuclei among cell clusters in snRNA-seq dataset. **C.** The module score of other cell-related genes (vasculature-related, guard cell, mesophyll cell, pavement cell, and subsidiary cell) in different cell clusters. **D.** Module scores for cell-type related genes projected on UMAP embedding.

